# Schrödinger’s mouse: human observation erases a drug-induced behavioral phenotype

**DOI:** 10.64898/2026.09.25.754328

**Authors:** Timothy A. Krausz, Thomas K. Roseberry, Gina Williams, David Tingley

## Abstract

Animal studies primarily rely on human monitoring for health and behavioral outcomes. The presence of a human, however, can induce stress and potentially alter experimental results. We sought to answer whether human observation alone is enough to change experimental outcomes and obscure the effect of interest. Our automated home-cage behavioral phenotyping system provided the ideal platform to test this hypothesis. Two cohorts received semaglutide (0.03 mg/kg, intraperitoneally) on separate consecutive days. The first cohort was recorded without inspection after dosing; the second underwent a brief health check. Recordings from the checked cohort on the preceding day, without injection or inspection, provided a matched baseline. A single brief check produced large, time-locked changes across the behavioral repertoire. The changes began before the cage door was opened, indicating a response to experimenter proximity. In addition to adding variance, the semaglutide-treated animals’ behavior during observation was more similar to the no-injection baseline than to the semaglutide/no-check reference, transiently erasing the drug phenotype. Brief human observation is therefore not a neutral measurement operation; over the minutes surrounding it, it is a perturbation in itself that is large enough to mask the drug effect it is meant to record.

## Introduction

Human involvement is a routine feature of laboratory animal research. Rodents are inspected for health and welfare (National Research Council, 2011), handled for dosing and experimental procedures (Gärtner et al., 1980; Diehl et al., 2001; Swan et al., 2024), and observed during behavioral testing (Irwin, 1968; Moy et al., 2009; Segelcke et al., 2023). Even studies with automated endpoints retain regular contact with animal-care personnel: the Guide for the Care and Use of Laboratory Animals recommends observation by trained personnel at least daily (National Research Council, 2011).

At the same time, research suggests that human experimenters can alter animal behavior, physiology, and experimental readouts. Rats are more likely to remain in opaque than clear cage compartments while experimenters are present (Cloutier and Newberry, 2010), and exposure to male experimenters or even male-associated odors induces stress-related analgesia in rodents (Sorge et al., 2014). Experimenter sex and olfactory cues were also found to modulate antidepressant-like responses to ketamine in mice (Georgiou et al., 2022). More recently, remotely delivered mechanical stimuli revealed that researcher presence altered mouse paw-withdrawal responses and produced sex differences that were not detected when testing was conducted remotely (Burdge et al., 2025). Human influence can also be seen at the neuron level: in bats, human identity and movement modulated hippocampal activity (Snyder et al., 2024). Together, these studies establish that human presence can influence a variety of experimental measurements and reported outcomes.

An important unresolved question is whether routine health checks transiently mask a drug-associated behavioral phenotype in the home cage. Mice can suppress pain responses during predator exposure or close contact with an unfamiliar male (Kavaliers and Colwell, 1994; Langford et al., 2011). Do mice mask their symptoms during health checks? This extends the problem of observer effects from particular assays and experimenter characteristics to an everyday component of animal care. A check could increase variability, shift behavior in a consistent direction, or make treated animals resemble untreated controls while they are being inspected. The latter would reduce the experimental signal available to the observer, with implications for interpreting behavioral signs of drug action or tolerability. Resolving these possibilities requires an uninterrupted record spanning the periods before, during, and after inspection, together with untreated and drug-treated reference profiles.

Our automated home-cage phenotyping platform provides this continuous view (Roseberry et al., 2026). The phenotype acquisition device (PAD) records behavior in the animals’ home environment and quantifies a broad repertoire of actions, movement, social interactions, and spatial features without requiring an observer to remain in the room. This allows a routine inspection to be examined as an event within an uninterrupted behavioral record and its effects to be evaluated against both untreated and drug-treated reference profiles.

Here, we used this platform to examine health checks in semaglutide-treated mice. A researcher opened each cage door and examined the mice for approximately one minute. Comparison with no-injection and semaglutide/no-check references revealed rapid changes across the behavioral repertoire, including increased walking and rotation, reduced sleep, and peaks in jumping and wall climbing before cage-door opening. Upon inspection, the checked animals’ behavioral profile shifted away from the semaglutide reference and became more similar to no injection before returning toward its pre-inspection state. These results show how a brief inspection can transiently obscure a drug-associated behavioral phenotype during the period in which the animals are being observed.

## Results

### Design

We used continuous video monitoring and behavioral quantification to characterize how behavior in semaglutide-treated mice changed around a routine health check. The PAD platform continuously records behavior across many cages and quantifies a broad repertoire of behavioral features (Roseberry et al., 2026; Fig. 1a,b). The analysis compared three conditions across two consecutive days: no injection without inspection, semaglutide without inspection, and semaglutide with inspection (0.03 mg/kg, intraperitoneally; Fig. 1c). Thirty cages provided the matched no-injection baseline on day 1 and the semaglutide-with-inspection condition on day 2. An independent cohort of 31 cages received semaglutide without inspection on day 1.

**Figure 1.**
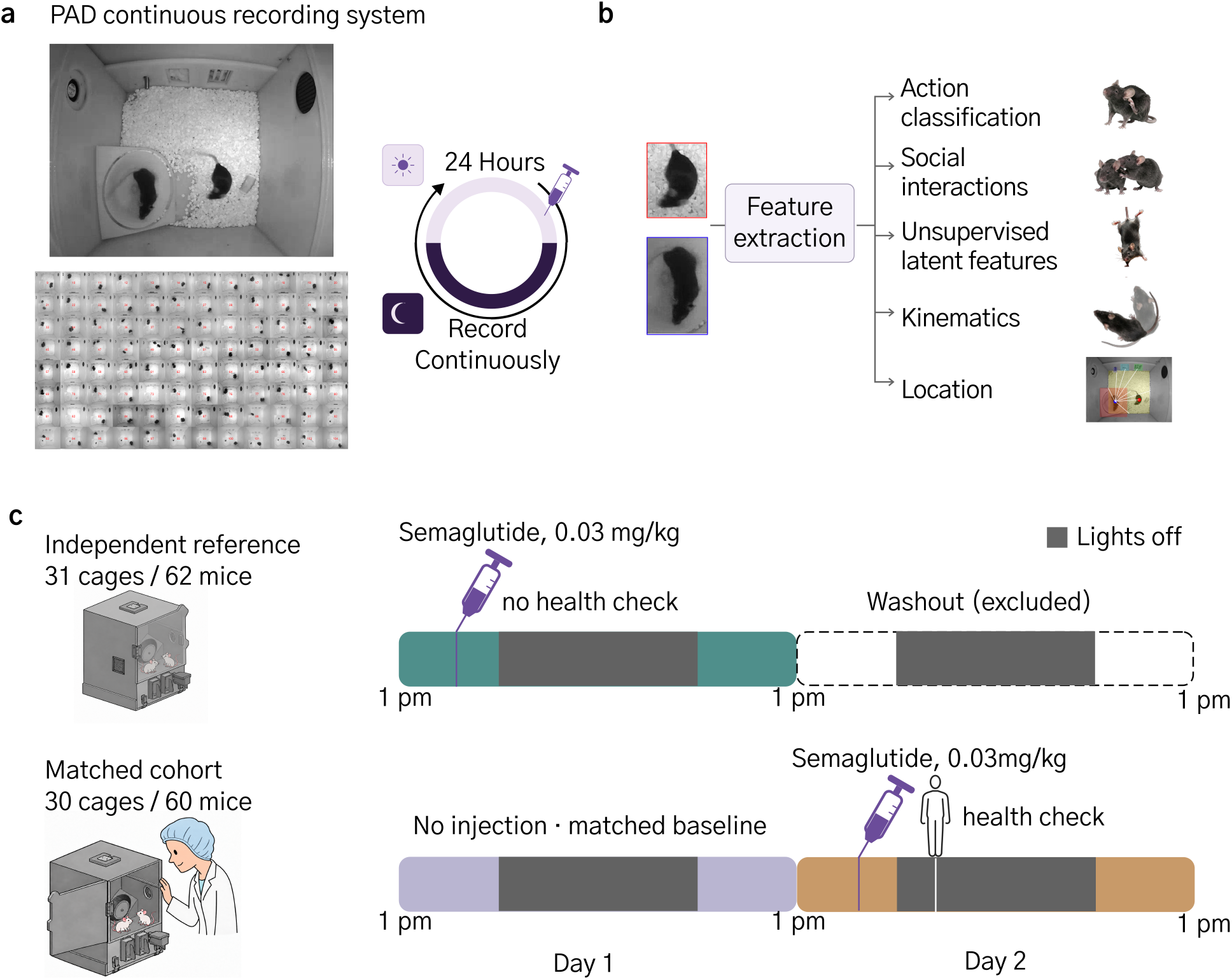
Experimental design. a) Top-down view of a phenotype acquisition device (PAD), 96 PADs recorded simultaneously and continuously 24/7. b) Video-derived feature extraction yields action classification, social interactions, unsupervised latent features, kinematics, and location (adapted from Roseberry et al., 2026). c) Two-cohort experimental timeline. An independent reference cohort (31 cages, 62 mice) provides the semaglutide-without-check condition. The matched cohort (30 cages, 60 mice) provides no-injection baseline and semaglutide-with-health-check conditions in the same cages across both days. Semaglutide dose: 0.03 mg/kg. Lavender, teal, and ochre denote no injection, semaglutide without check, and semaglutide with check, respectively; gray shading denotes lights-off. Experimental procedures are described in Methods.

The two semaglutide conditions were recorded on separate nights to avoid exposing the no-check cohort to an experimenter checking other cages in the same room. Mice were already habituated to daily handling. Checks were timed to coincide with the period when the behavioral effects of semaglutide were expected to be greatest. Each check involved opening the cage door and examining the mice for at most one minute. Time zero in the event-aligned analyses denotes cage-door opening or matched reference times for control groups.

### Inspection coincides with rapid changes across the behavioral repertoire

Examining behavioral dynamics over the course of each experimental day revealed a transient increase in walking for the checked group around the time of inspection (Fig. 2a). Event alignment resolved an abrupt increase near time zero that diminished over the subsequent minutes (Fig. 2b). Between the pre-inspection window (*−* 5 to *−*4 min) and the first minute after cage-door opening, median walking proportion increased from 0 to 0.0419 (95% confidence interval [CI], 0.0249–0.0536; adjusted P = 0.00024; Fig. 2b). Right rotation amount also increased from a median of 0 to 3.56 (95% CI, 2.91–5.23; adjusted P = 0.00024; Fig. 2c), while sleep proportion decreased from 0.794 to 0.155 (post-inspection 95% CI, 0.084–0.287; adjusted P = 0.00024; Fig. 2d). No corresponding within-cohort changes were detected in either unchecked reference after correction across the original 12 paired tests (all adjusted P *≥* 0.248).

**Figure 2.**
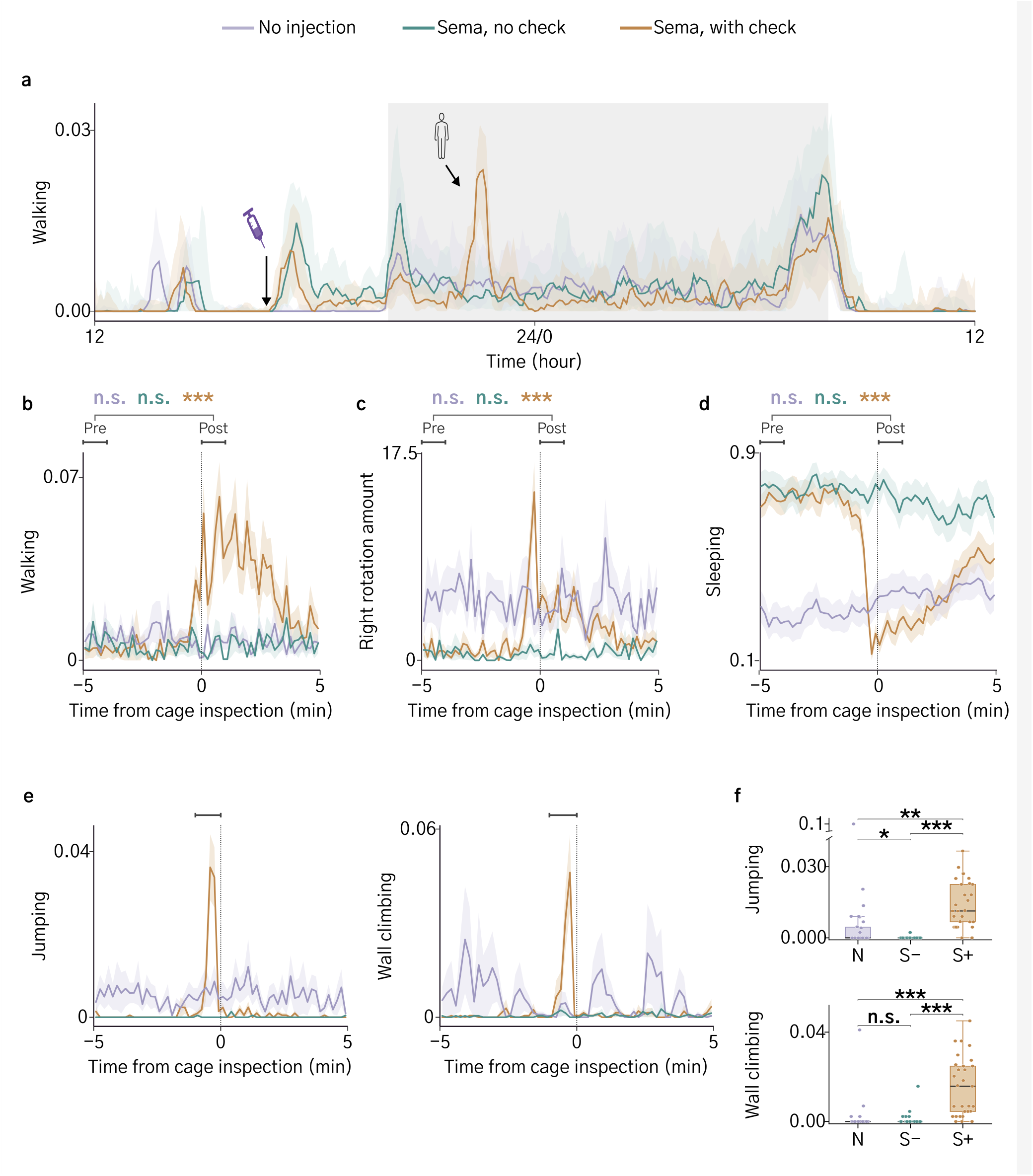
Human presence changes mouse behavior. Lavender, no injection (N); teal, semaglutide without check (S *−*); ochre, semaglutide with check (S+). a) Daily walking profiles over the course of each experiment; lines and ribbons show cohort medians and interquartile ranges. Syringe and human icons denote dosing and cage inspection, respectively; gray shading denotes lights-off. b, c, d, and e) Event-aligned walking, right rotation, sleeping, and jumping/wall-climbing traces, respectively, displayed from *−* 5 to +5 min; lines and ribbons show means *±* SEM. Time zero denotes cage-door opening, with corresponding reference times for unchecked cohorts. b–d) Colored symbols above the bracket linking pre [*−* 5, *−* 4) and post [0,1) min windows indicate within-cohort pre/post tests, in N/S *−* /S+ order; these are window-level tests, not tests at individual time points. b and d use n = 30/31/29 pre and 30/30/28 post; c uses 30/31/30 pre and 30/30/29 post. Paired test sample sizes in N/S *−* /S+ order are 30/30/28 for b and d and 30/30/29 for c. f) Jumping and wall climbing during [*−*1,0) min (n = 30/30/29). Boxes show medians and quartiles, whiskers extend to the most extreme observations within 1.5 IQR of the quartiles, and points represent cages. Colored symbols in b, c, and d indicate within-cohort pre/post comparisons; brackets in f indicate condition comparisons. S+ adjusted P= 0.00024 in b, c, and d; all N and S *−* pre/post comparisons are nonsignificant. ^*^ P < 0.05; ^**^ P < 0.01; ^***^ P < 0.001; n.s., P *≥*0.05 (all adjusted). Processing and statistical procedures are described in Methods. Within-window adjusted P values are listed in N versus S *−*, N versus S+, and S *−* versus S+ order: b: pre (0.29256, 0.01944, 1); post (0.13678, 0.00036, 0.00036). c: pre (0.00084, 0.00408, 0.79454); post (0.0092, 0.72364, 0.00024). d: pre (0.00024,0.00024, 0.17008); post (0.00032, 0.17008, 0.00024). f: jumping (0.011, 0.00264, 0.00036); wall climbing (1, 0.00036,0.00036).

Other behaviors changed before the door was even opened. Jumping and wall climbing showed brief peaks immediately before cage-door opening (Fig. 2e). During the *−*1 to 0 min window, jumping was higher in checked cages than in both the no-injection and semaglutide/no-check references (adjusted P = 0.00264 and 0.00036, respectively). Wall climbing was also higher than in both references (adjusted P = 0.00036 for each comparison; Fig. 2f).

### Inspection erases the drug phenotype during observation

Feature dynamics across all extracted behavioral features revealed broad differences between the semaglutide and no-dose controls. Prior to inspection, the checked semaglutide group appeared similar to the semaglutide control. Around the time of cage inspection, however, the behavioral profile changed across features to resemble the no-injection reference (Fig. 3a).

**Figure 3.**
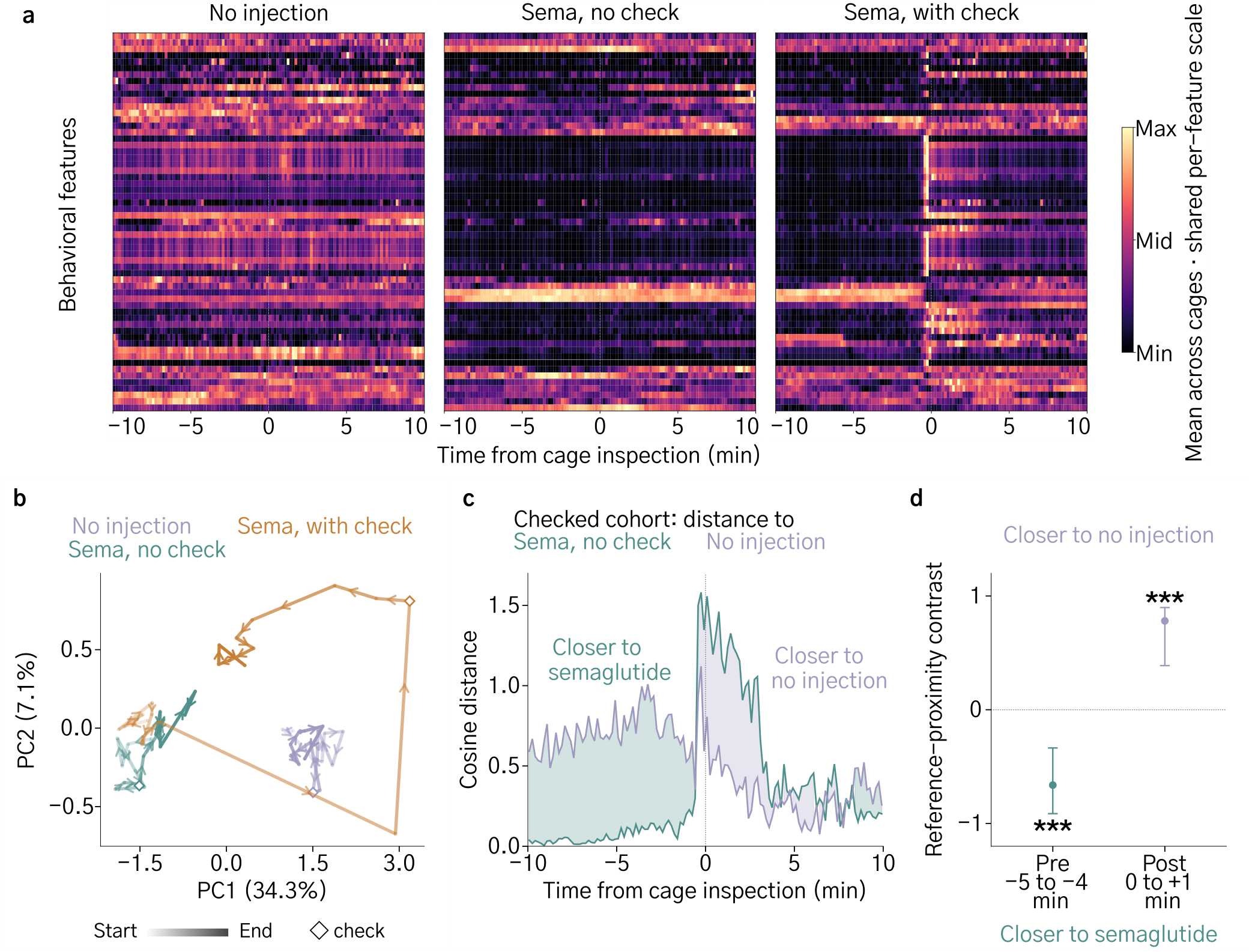
Human observation erases the drug phenotype. All panels use the same 59 behavioral features. a) Cohort-mean feature dynamics; each row uses a shared color scale across conditions, from its minimum to maximum. b) Mean cohort trajectories in principal-component space. Dots show successive one-minute bins; arrows on every segment indicate temporal order. Trajectory opacity increases from 18% at the start to 100% at the end; arrow opacity is kept at a minimum of 55% for visibility. Open diamonds mark the [0,1) min bin. PC1 and PC2 explain 34.3% and 7.1% of variance, respectively. c) Cosine distance from the checked cohort to the semaglutide/no-check reference (teal line) and no-injection reference (lavender line). Shading identifies the closer reference: lavender, no injection; teal, semaglutide. d) Reference-proximity contrast (distance to semaglutide minus distance to no injection) during pre [*−*5, *−* 4) and post [0,1) min. Positive values indicate greater similarity to no injection; negative values indicate greater similarity to semaglutide. Points show window means and error bars show 95% confidence intervals (30 matched cage pairs and 31 independent reference cages). The pre-event contrast was *−* 0.662 (95% CI, *−*0.913 to *−*0.336), and the post-event contrast was 0.781 (95% CI, 0.388 to 0.898); adjusted P = 0.0002 for both comparisons against zero (^***^ P < 0.001). Time zero denotes cage-door opening. Feature processing, PCA, distance calculations, and statistical procedures are described in Methods.

To test whether cage inspection made behavior less semaglutide-like, we first explored dynamics in a low-dimensional space. PCA of the features revealed a pronounced excursion of the checked cohort’s mean trajectory away from the semaglutide control’s behavioral space, toward and around the no-dose space at cage-inspection time, followed by movement back toward the semaglutide control region (Fig. 3b).

To quantify reference proximity in the full feature space, we compared cosine distances from the checked cohort to the two unchecked references (Fig. 3c). Before inspection, the checked cohort’s behavioral features were more similar to those of the no-check semaglutide group. In the first minute after cage-door opening, however, its profile was closer to the no-injection reference. The mean proximity contrast, defined as distance to semaglutide control minus distance to no injection, changed from *−* 0.662 (95% cage-bootstrap CI, *−* 0.913 to *−*0.336) in the pre-inspection window to 0.781 (95% CI, 0.388–0.898) after inspection (Fig. 3d). Each contrast differed from zero after correction across the two window tests (adjusted P = 0.0002 for both). Across the day, the drug-associated behavioral profile varied with circadian phase and estimated plasma exposure, with greater separation from no injection after lights-off and a transient disruption during human inspection (Supplementary Fig. 1a). Extended event alignment showed that more than an hour elapsed before the checked cohort stably returned toward the uninspected semaglutide profile (Supplementary Fig. 1b). Thus, inspection was associated with a transient loss of the semaglutide behavioral phenotype, and this loss was strongest during the inspection itself.

## Discussion

A brief health check coincided with a coordinated shift in semaglutide-treated mice toward the no-injection behavioral profile. This extends evidence that experimenter cues alter pain responses and pharmacological effects (Sorge et al., 2014; Georgiou et al., 2022; Burdge et al., 2025) to routine observation in the home cage. Continuous recording revealed that the drug-associated phenotype was least apparent during the period when the animals were being inspected, creating a bias in assessments.

While the health checks included brief contact, the mere presence of the experimenter was enough to change behavior. Indeed, numerous behaviors increased before the cage door was ever opened. Whether jumping and wall climbing were escape behaviors induced by stress, exploratory behaviors to better smell the human cues in the air, or something else altogether will require further testing.

The findings also motivate closer examination of how welfare is assessed. Automated home-cage monitoring has detected reduced locomotion, climbing, and rearing in lipopolysaccharide-treated mice, demonstrating sensitivity to sickness-related behavioral changes (Hasriadi et al., 2021). If inspection transiently suppresses such signs, intermittent human checks systematically underestimate their persistence.

These findings argue for designing animal experiments around continuous automated monitoring and removing routine in-person checks wherever validated remote measures can meet welfare requirements. Automated systems quantify the timing, duration, and coordination of behaviors across many cages with a resolution and consistency far beyond what is possible from a brief visual inspection (Hasriadi et al., 2021; Roseberry et al., 2026). Combined with evidence that inspection can mask the phenotype being assessed, this makes minimizing human intervention a priority for both measurement quality and welfare surveillance. Indeed, replacing routine in-person health checks with validated remote monitoring would advance refinement within the FDA-supported 3Rs by minimizing potentially stressful human disturbance while maintaining welfare surveillance (U.S. Food and Drug Administration, 2022). Human assessment should be targeted to clinical need and essential care, supported by automated alerts and veterinary oversight. The aim should be to measure animals in their undisturbed state and intervene when necessary, rather than routinely perturbing the state we seek to assess.

This study also underscores the importance of animal assay design for human translation. A biologically informative mouse model cannot yield useful predictions if measurement suppresses or distorts the relevant signals. Indeed, translational limitations are evident across widely used behavioral assays. A meta-analysis of common mouse anxiety tests, including the elevated plus maze and open field, found that only two of 17 outcome measures reliably detected the effects of drugs used to treat anxiety in humans (Rosso et al., 2022). Similarly, a retrospective analysis of pharmaceutical compounds tested in the forced swim assay found limited success in predicting antidepressant efficacy in humans (Trunnell and Carvalho, 2021). Our study exposes an additional measurement problem in one of the most routine practices in animal research: the daily health check. Before attributing poor translation to the animal model, we must establish that our assays are optimized to detect the biology of interest rather than the response to being measured. Improving translation therefore requires better signal acquisition, beginning with how we observe the animals each day.

## Methods

### Animals and experimental design

The analysis used recordings from two consecutive experiments conducted on 22–24 June 2026 in a continuously monitored home-cage platform. Selected cages housed two mice of the same sex, and each cage was treated as the sampling unit. Animals were C57BL/6J mice from Jackson Laboratories or C57BL/6NTac mice from Taconic and were approximately six months old at recording. Animal care, ethical oversight, husbandry, and PAD design followed the platform procedures described by Roseberry et al. (2026). The experimental procedures in this study were approved by the same external Institutional Animal Care and Use Committee described in Roseberry et al. (2026). Mice were housed in same-sex pairs in PADs equipped with running wheels, under a 12 h light/12 h dark cycle (lights on at 08:00 and off at 20:00), with ad libitum water, high-fat diet (Research Diets D12492i), and regular chow (LabDiet PicoLab 5053) in separate hoppers. The platform protocol included 10 days of PAD habituation and at least seven days of behavioral observation before experimental enrollment. Mice were handled daily for a minimum of 10 days during habituation by both male and female researchers, providing repeated exposure to humans before the experimental checks. The three-condition analysis used 30 cages observed without injection or inspection on day 1 and after semaglutide administration and inspection on day 2, together with 31 independent cages treated with semaglutide without inspection on day 1. Semaglutide was administered intraperitoneally at 0.03 mg/kg. No-dose animals received neither drug nor vehicle injection. On the no-check day, experimenters stayed out of the testing room after dosing. The day-2 washout period of the independently treated cohort was excluded from these comparisons. PADs were randomly assigned to experimental groups, and the experimenter conducting cage checks was blinded to group assignment. Semaglutide/no-check and semaglutide/check recordings were obtained on separate nights to avoid exposing no-check cages to an experimenter checking other cages in the same room. This design prevented concurrent exposure of the no-check group but left check status linked to experiment day.

The matched cohort comprised 20 female and 10 male cages, equally divided between the two animal sources within each sex. The analyzed control semaglutide cohort comprised 31 cages: 17 female and 14 male. Values from the two mice were averaged within each cage for individually resolved features; mice sharing a cage were not treated as independent replicates. Cage and box identifiers were used jointly to preserve matching across days.

### Event timing and reference alignment

On the second night, the experimenter performed sequential health checks only on boxes containing mice that had received semaglutide that day. Boxes containing mice that had not been injected that day were not opened or checked. Checks were timed to coincide with the greatest expected behavioral effects of semaglutide, based on previous internal observations and expected pharmacokinetics. The PK overlay in Supplemental Figure 1 is an illustrative model, normalized to peak plasma concentration. It is consistent with an assumed 8 h elimination half-life, informed by the mouse terminal half-life reported by the European Medicines Agency (2017, section 2.3.3, p. 24). Intraperitoneal absorption kinetics and plasma concentrations were not measured in this study. Under a monoexponential post-peak approximation, *C*(24 h)/*C*_max_ = 2^*−*(24*−*1.3)/8^ *≈* 0.14; this is a model-derived value, not an observed concentration. For each check, a male researcher opened the cage door and examined the mice for approximately one minute. This included brief and gentle skin contact to check for responsiveness. Time zero was each cage’s door-opening time, recorded by a QR scan. The 30 event times ranged from 21:56:54 to 22:24:17 on 23 June 2026. Negative times therefore preceded cage-door opening; the onset of the experimenter’s presence or approach was not separately timestamped.

Each matched no-injection cage was aligned to its own day-2 event time minus one calendar day. Independent semaglutide/no-check cages were assigned the median event clock time of checked cages in the same sex-by-source stratum. Acute analyses covered [*−*10, +10) min relative to these times. Bins were half-open and labeled by their centers, giving 120 ten-second bins centered from *−*595 to +595 s. Reference times did not indicate an actual inspection in the unchecked cohorts.

### Video-derived behavioral features and temporal summaries

Behavioral features were obtained using the video-processing pipeline described by Roseberry et al. (2026), including pose-based kinematics and spatial measures, social features, and per-frame action classifications. Action outcomes were calculated as the proportion of classified frames assigned to the corresponding action and averaged across the two mice within each cage. Right rotation amount was retained as the pipeline’s continuous, pose-derived angular rotation feature rather than a behavioral proportion. Acute analyses used processed, unbinned feature and action files, aggregated into ten-second bins.

Event-aligned traces show means and standard errors across available cages in each ten-second bin. Event-aligned analyses covered *−* 10 to +10 min; Figure 2 displays the central *−*5 to +5 min interval using the original ten-second bins. For the daily walking profile, a centered four-bin rolling mean, requiring at least two available bins, was applied to five-minute summary data within each cage before calculating cohort medians and interquartile ranges. Observed zeros were retained, and missing observations were excluded from each available-case summary rather than replaced with zero.

### Cage-level window summaries and statistical comparisons

Walking, right rotation amount, and sleep were summarized in pre [*−*5, *−* 4) and post [0, 1) min windows. Jumping and wall climbing were summarized separately over [*−*1, 0) min. A cage contributed the mean of its six ten-second bin values only when all six were finite. For walking and sleep, sample sizes in no-injection, semaglutide/no-check, and semaglutide/check order were 30/31/29 before and 30/30/28 after inspection. Rotation sample sizes were 30/31/30 before and 30/30/29 after inspection. Jumping and climbing each used 30/30/29 cages. Marginal cohort medians were reported for each window, with pointwise 95% percentile confidence intervals from 99,999 cage-bootstrap draws. Sparse outcomes could yield zero-width intervals. Box plots show medians, quartiles, whiskers extending to the most extreme values within 1.5 interquartile ranges, and individual cage values.

Within each cohort, pre/post comparisons used two-sided paired Wilcoxon signed-rank tests with Pratt handling of zero differences (Pratt, 1959) and permutation-based P values from 99,999 resamples. Complete pairs were matched by box and cage identifiers. Walking, head shaking, and sleep used 30/30/28 pairs, and rotation used 30/30/29 pairs, in the cohort order above. Differences were rounded to 12 decimal places to avoid numerical rank artifacts; all-zero paired differences received P = 1. Holm correction (Holm, 1979) retained the original family of 12 pre/post tests across four features and three cohorts, including the three head-shaking tests omitted from Figure 2. These tests assessed paired changes; the reported medians and confidence intervals used each window’s complete marginal sample.

Within-window condition comparisons used matched-cage Wilcoxon-Pratt tests for no injection versus semaglutide/check and Mann–Whitney U tests for comparisons involving independent semaglutide/no-check cages. Tests were two-sided, with 99,999 permutation resamples and fixed random seeds. Degenerate comparisons with identical samples were assigned P = 1. The original Holm families were retained as plots were revised: 18 comparisons across digging, walking, jumping, and climbing, and 12 across rotation and sleep. Digging comparisons remained in their original correction family although the corresponding panels are not displayed. Significance was evaluated at adjusted P < 0.05, and confidence intervals were not adjusted for multiplicity.

### Multivariate heatmaps and spatial features

Heatmaps contained 51 behavioral features, three camera-edge features, and five spatial occupancy principal components. Each cell was the mean of available cage values within a ten-second bin. For each feature, the minimum and maximum across all three cohort-mean traces and all time bins defined a common color scale. This transformation affected color mapping only and was not within-cohort normalization or baseline subtraction.

Spatial occupancy was reconstructed from individual bounding-box centers on a 20 × 20 grid spanning the 640 × 480-pixel camera frame. Occupancy fractions were averaged across available mice within each cage and bin. Five principal components were fitted to pooled cage-bin occupancy profiles from all three conditions using centering without standardization or whitening, so all groups shared the same axes. These components explained approximately 24.5% of occupancy variance. Camera-edge features were cached histogram-density measures of distance to image boundaries, not calibrated distances to physical cage walls. The invariant edge feature was omitted.

### Principal-component trajectories

The trajectory analysis used the same 59 behavioral features as the heatmaps. Each observation was a cage-minute. A feature’s minute mean required six finite ten-second bins. Cage-minutes with fewer than 90% of features observed were excluded; remaining missing entries were imputed with feature medians from the pooled prior-day (day 0, prior to the experiment) reference cohorts. Feature means and population standard deviations from the imputed reference observations were used to standardize all conditions. An inverse hyperbolic sine transformation was then applied to standardized values, without a second standardization.

PCA was fitted jointly to all 1,759 retained cage-minute observations, without condition labels, and the same fitted coordinates were used throughout. Trajectories show arithmetic means of available cage coordinates in 20 consecutive one-minute bins. Arrows indicate temporal order, increasing opacity indicates progression from the first to the last bin, and open diamonds mark the [0, 1) min bin. PC1 and PC2 explained 34.3% and 7.1% of variance, respectively.

### Cosine distance and reference-proximity inference

Cosine analyses used the same 59-feature set at ten-second resolution, without the inverse hyperbolic sine transformation. A mean and sample standard deviation were estimated for each feature from both prior-day reference cohorts pooled across the full 20-minute interval. This common scale was applied to all three conditions. At each bin, cohort-median standardized feature vectors were compared using cosine distance, 1 *−* (x·y)/(‖x‖‖y‖), using a common finite-feature mask across all three cohort vectors. Missing observations were not imputed in this analysis. The proximity contrast was the distance from the checked cohort to semaglutide/no-check minus its distance to no injection. Positive values indicated greater proximity to no injection, and negative values indicated greater proximity to semaglutide/no-check. Each window estimate was the mean of six ten-second contrasts calculated before temporal averaging.

Uncertainty was estimated with 9,999 cage-bootstrap resamples, stratified by sex and animal source. The 30 matched checked/no-injection cages were resampled together, while the 31 independent semaglutide cages were resampled separately. Entire cage profiles were sampled to preserve temporal dependence. Reference scaling, cohort medians, distances, and window contrasts were recalculated in each resample. The shared occupancy-PC basis used in all panels was held fixed during resampling. All resamples yielded finite estimates. Error bars indicate marginal percentile 95% confidence intervals. Approximate two-sided tests against zero compared the absolute observed contrast with absolute bootstrap deviations centered on that contrast, using the finite-resample correction (extreme count + 1)/(9,999 + 1). Holm correction covered the two window tests. The adjusted P value of 0.0002 reflects finite Monte Carlo resolution and is not an exact tail probability.

---

**Supplemental Figure 1.**
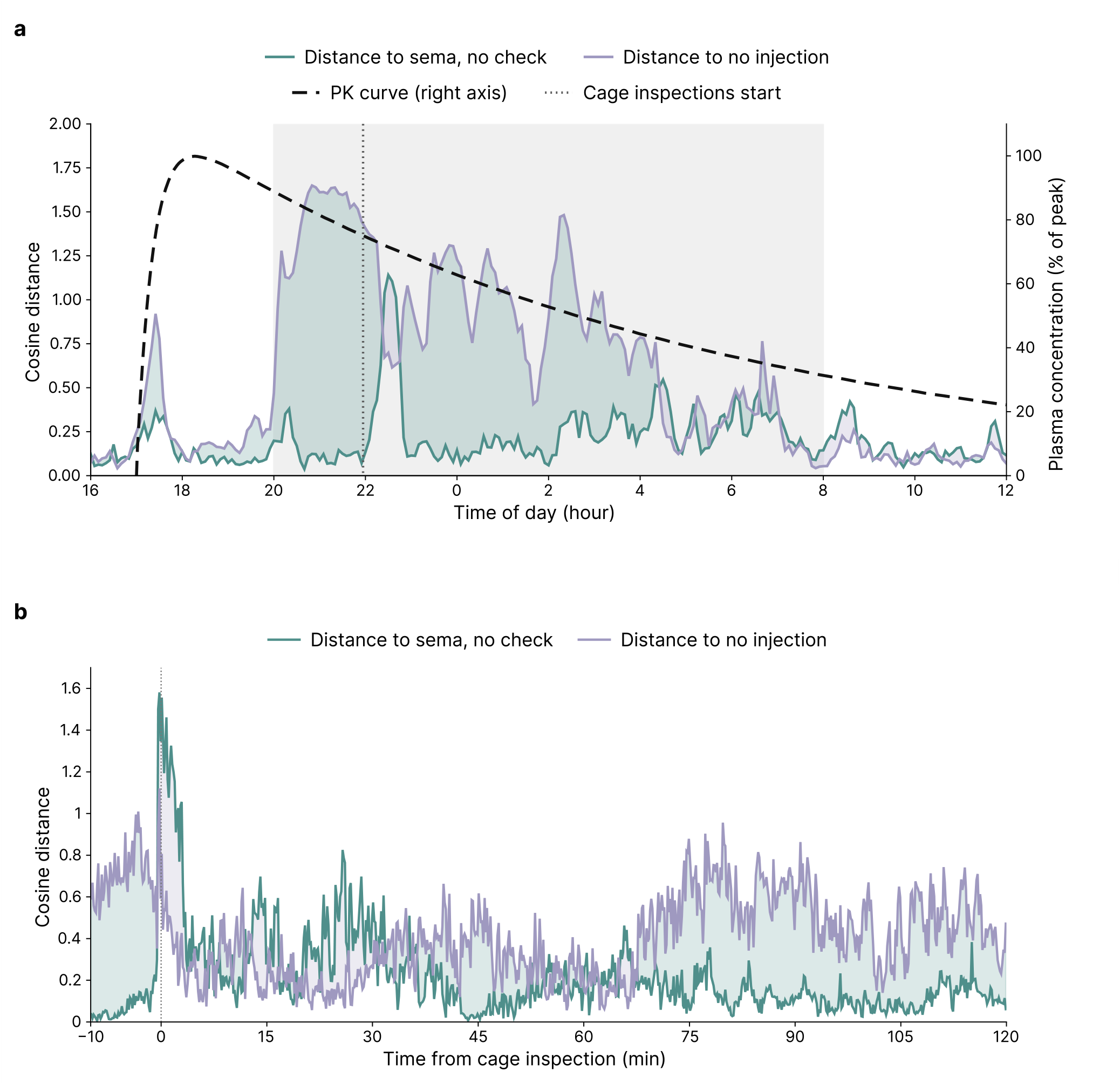
Circadian, pharmacokinetic, and cage-inspection effects on behavioral similarity. (a) Cosine distances compare the semaglutide + cage-inspection cohort with semaglutide without inspection (teal) and no injection (lavender); lower values indicate more similar behavioral profiles. Native 5-minute cage summaries were smoothed with a centered 20-minute rolling mean. Cohort-median vectors of the same behavioral features as in Figure 3 were standardized using the pooled reference cohorts’ full-day mean and sample SD. Distances were calculated at matching Pacific clock times; colored fill indicates the closer reference. Gray shading marks the lights-off period. The black dashed curve shows an illustrative PK model normalized to peak plasma concentration (right y-axis); model assumptions are described in Methods. (b) The corresponding cage-inspection-aligned comparison from *−*10 to +120 minutes (extended from Figure 3c). The dotted line marks cage inspection.

## Notes

### Competing Interest Statement

All authors are employees of Olio Labs.

